# Comparison and evaluation of data-driven protein stability prediction models

**DOI:** 10.1101/2022.03.14.483859

**Authors:** Jennifer A. Csicsery-Ronay, Alexander Zaitzeff, Jedediah M. Singer

## Abstract

Predicting protein stability is important to protein engineering yet poses unsolved challenges. Computational costs associated with physics-based models, and the limited amount of data available to support data-driven models, have left stability prediction behind the prediction of structure. New data and advancements in modeling approaches now afford greater opportunities to solve this challenge. We evaluate a set of data-driven prediction models using a large, newly published dataset of various synthetic proteins and their experimental stability data. We test the models in two separate tasks, exercising extrapolation to new protein classes and prediction of the effects on stability of small mutations. Small convolutional neural networks trained from scratch on stability data and large protein embedding models passed through simple downstream models trained on stability data are both able to predict stability comparably well. The largest of the embedding models yields the best performance in all tasks and metrics. We also explored the marginal performance gains seen with two ensemble models.

## 1 Introduction

A protein’s stability—the likelihood it assumes its folded configuration—is an essential property that interacts with its structure, function, solubility, and durability [11]. Additions, deletions, and alterations of single amino acids can change not only the folded structure but also its stability [13]. Predicting a change in a protein’s stability under the effect of a mutation can guide engineering in medically significant domains, such as diagnoses, therapeutics, and vaccine development [3, 7]. However, stability prediction remains challenging.

Physics-based software like Rosetta drives computational protein design and is responsible for many stable synthetic proteins [9, 15]. Despite recent progress, however, statistical and physics-based methods for stability prediction remain computationally expensive and struggle to model and predict the quantities and interactions governing protein stability [8, 10]. Meanwhile, advancements in data-driven methods such as high throughput stability measurement, deep sequencing, and machine learning are beginning to mitigate issues of computational cost and prediction accuracy [1, 7, 12, 16, 18].

Here we present a rigorous comparison and evaluation of several data-driven stability prediction models using 112,455 synthetic proteins from a recently published dataset [16] of experimental stability data. We evaluate performance on held-out protein classes to test models’ ability to extrapolate new kinds of protein designs. We also further the work of [12] and evaluate models’ predictions on single-site mutations to test their ability to predict small changes in the stability landscape. Models tested include a convolutional neural network (CNN) model from [16] and three simplified variants of this model, as well as gradient boosting models built using three recently-published protein embeddings inspired by natural language processing: (i) the Evolutionary Scale Model (ESM, [14]), (ii) ProtBERT [6], and (iii) UniRep [1]. We use XGBoost [4] to construct these gradient boosting models and test both default and optimized parameters. Finally, we use two linear blending techniques to assemble ensemble models and evaluate their performance against their constituents.

## 2 Methods

### 2.1 Data

Models were trained on data for 112,455 proteins, reported in [16]—a superset of data from [15]. We restrict ourselves to expert-designed proteins with explicitly intended topologies, as we use design methodology and intended topology to test model extrapolation. The data consist of primary sequences for small synthetic proteins, as well as experimentally measured stability scores [15]. We evaluate models on subsets of these data as well as near-comprehensive single-site mutations for seventeen proteins (three natural and fourteen synthetic), also originally reported in [15].

### 2.2 Stability Prediction Tasks

We evaluate each model on two stability prediction tasks: (i) predicting stability for members of a single class (defined by secondary structure topology and design method) held out from the training set, and (ii) predicting stability for near-comprehensive single-site mutations of known proteins. For the held-out class approach, We use each class in turn as the test set and train each of the models on the other 26 topologies. To verify that the proteins in each topology were distinct, [16] used BLAST to identify the nearest homolog of each protein from the other 26 classes. The results yielded low maximum bit scores and maximum percent identities (mean ± SD of 34.5 ± 5.1 and 47.0 ± 8.3 respectively), suggesting considerable differences among each topology class [2, 16]. This task evaluates the models’ ability to generalize or extrapolate to new kinds of proteins and could be important for downselecting proteins to test from a larger library of designs. For the single-site mutation predictions, we follow the approach of [12], testing on a set constructed from 17 1-Hamming distance single-site mutations from [15]. Note that for our experiments we used a larger set of training data than in [12]. This task evaluates models’ ability to predict the effects of small mutations on protein stability and could be important for making incremental improvements to the stability of proteins with desired properties.

### 2.3 Metrics

For the held-out topology task, we evaluated the performance of each model using the *R*^2^ goodness of fit score (i.e. the coefficient of determination), root mean squared error (RMSE), average precision (AP), and the area under the receiver operating characteristic curve (ROC AUC). *R*^2^ and RMSE directly compare raw stability values to predicted stability scores. *R*^2^ tells us what fraction of the variance in the data the model captures beyond simply predicting the mean. RMSE tells us how close, in stability score units, model predictions are to observed values, with more weight given to large errors. We prefer *R*^2^ because different protein classes have different ranges of stability, some of which are quite small relative to the noise in the stability score assay; RMSE can therefore be misleading to compare across held-out classes. AP and ROC AUC depend on a definition of “stable”, and measure the models’ ability to rank stable proteins more highly than unstable ones. Average precision, essentially the area under the precision/recall curve, encapsulates the tradeoff between false positives and false negatives. The receiver operating curve measures true positives against false positives. AP tends to be more robust to class imbalance than ROC AUC, but ROC AUC sees wider use. We follow the convention adopted in [15] and call a protein stable if its stability score is at least 1.0.

To evaluate performance in the single-site mutation prediction task we chose Spearman’s rank correlation (*ρ*), as well as *R*^2^ and RMSE. Spearman’s *ρ* assesses the degree to which predicted stability scores sort in the same order as observed stability scores. This task asks how well a model can predict the effect on stability of small changes relative to a baseline. To guide stabilizing single-site mutations, it is sufficient to sort stability scores successfully, making Spearman’s *ρ* a relevant metric. Average precision and the area under the ROC curve are less relevant to this task; single-site mutations tend to change stability by a fairly small amount, so in general either most mutants will be stable (stability score ≥ 1) or most will be unstable.

### 2.4 Stability Prediction Algorithms

We evaluated the ability of two types of models to predict protein stability. One type consisted of convolutional neural network models—CNNs—trained on our stability data to directly predict stability scores. The other type was protein sequence “embeddings”, large models that attempt to convert an amino acid sequence into a fixed-dimensional vector that represents a variety of properties of the encoded protein. For these embedding models, we used gradient boosting to relate these embedding vectors to stability scores.

We considered four variants of CNN models. The most complex was the previously-published Evaluator Model (EM) from [16]. This model expands upon a typical CNN architecture by jointly predicting stability score and protein secondary structure, and learning from synthetic and natural proteins. We also considered three simpler variants of the EM: (i) trained with no natural proteins or secondary structure prediction, (ii) trained with natural proteins but not secondary structure prediction, and (iii) trained with secondary structure prediction but not natural proteins.

Gradient boosting models were trained with XGBoost (XGB) [4] with the following parameters chosen via GridSearchCV: n_estimators=1000, colsample_bytree=0.5, learning_rate=0.01, and max_depth=8. These parameters optimized *R*^2^, and were chosen from n_estimators ∈ {1, 100, 1000}, colsample_bytree ∈ {0.3, 0.5, 0.7}, learning_rate ∈ {0.001, 0.01, 0.1}, and max_depth ∈ {4, 6, 8}. The hyperparameters for each gradient boosting model were optimized independently, but all three models performed best with the same set of hyperparameters. The training objective for the gradient boosting models was the minimization of mean squared error.

We also evaluated gradient boosting models trained using the default XGB parameters: 50 estimators, 1 column to be randomly sampled per tree, a learning rate of 0.3, and a maximum tree depth of 6. We apply XGB to features generated by extracting embeddings from sequence data by the ProtBERT model [6], the evolutionary scale model (ESM) [14], and the UniRep model [1]. Each of these models was trained in a self-supervised learning task; we used the published, pre-trained versions. Each yields a fixed-length vector for a given protein, which encodes some degree of biological information. We go into more detail on each of the embedding models below.

The ProtBERT model [6] uses the Bidirectional Encoder Representations from Transformers (BERT) [5] architecture applied to proteins. BERT is built on the powerful Transformer architecture [17] from natural language processing and is trained to predict masked or “missing” words from sentences; ProtBERT treats protein primary sequences as sentences, and amino acids as words, and learns to predict masked amino acids. The ProtBERT model was trained on the UniRef100 database and yields a 1024-feature vector per protein, implicitly encoding structural and functional information.

The Evolutionary Scale Model (ESM, [14]) is similar to ProtBERT in that it uses a series of deep Transformers [5] to learn the statistics and correlation of amino acid sequences and their contexts. It is a larger model, 650 million parameters to ProtBERT’s 420 million, and is trained on a set of 250 million diverse proteins from the UniParc, UniRef50, and UniRef100 databases [14]. This process results in a 1280-feature vector for each protein sequence carrying information about biological, biochemical, and structural protein properties. Like ProtBERT, the model was trained to predict masked amino acids from the primary sequences of natural proteins.

The UniRep model [1] implements a multiplicative Long-Short-Term-Memory recurrent neural network trained on 24 million sequences from the UniRef50 database, without the use of structural or evolutionary data. This model processes the input sequence one amino acid at a time and learns to predict at each point the next amino acid based on the sequence thus far. We used the UniRep Fusion variant, which yields 5700-dimensional vectors (concatenations of mean hidden state, final hidden state, and final cell state) containing approximations of encoded functional, structural, and evolutionary information.

For each XGBoost model, We randomly selected 10% of the training data to use for validation, with early-stopping governed by the validation loss. For each CNN model 10k was randomly selected as validation data with custom loss functions calculating the difference among a combination of the model’s stability, natural comparator, and dssp loss given the variation of the model. Both the CNN and XGBoost models have an early stopping value of 5 iterations for which the next decrease in loss is calculated. Each model was trained three times, with the final prediction for each model being the mean of the predictions from the three training runs.

Finally, we consider two linear regression ensemble models. The first used a “blending” architecture for both the held-out topology and single-site mutation tasks. Blending is an ensemble learning technique that helps to improve upon the accuracy of results from a set of models by learning how best to combine their predictions. Each ensemble model consists of the EM and the three tuned gradient boosting models. We built one ensemble model for each of the 27 held-out protein classes, and one to predict the stability of single-site mutations. Blending model coefficients optimized the mean squared error over the data sets used to fit them. The second ensemble approach was to simply take the mean predictions of the four models.

## 3 Results and Discussion

### 3.1 Model Performance

#### 3.1.1 Held-out Topology Task

The ESM gradient boosting model matched or outperformed all other individual models across all metrics for this task, with a mean *R*^2^ of 0.22 across the 27 protein classes. When compared between the default XGBoost parameters and the GridSearchCV results, the ESM gradient boosting model outperformed both the ProtBERT and UniRep gradient boosting models. Given the ESM model had been trained on the largest dataset (86 billion amino acids across 250 million diverse proteins), with their model trained with 650 million parameters [14], this suggests larger datasets could increase model performance.

The blending model, which fit a linear regression to combine the predictions of the EM and the three tuned embedding models, yielded only marginal improvements over the ESM model. This may reflect that the other models are failing to represent very much information that is not represented by the ESM. However, we note that a naïve ensemble that simply uses the mean of the four constituent models slightly outperforms the blending model on the preferred metrics (*R*^2^ and average precision). So it may instead be that coefficients that are optimal for the linear fit over proteins from classes familiar to the constituent models are inappropriate for extrapolating to new protein classes.

#### 3.1.2 Single-Site Mutation Task

The ESM gradient boosting model showed the highest ranking success when assessing how well the landscape of stability was learned (as suggested by [12]) (Spearman *ρ* = 0.73). Both the EM and ESM gradient boosting model tied in performance (*R*^2^ = 0.5). Scores were calculated from the mean of three runs per model.

We note that the UniRep gradient boosting model (*ρ* = 0.67) did not fare as well as was reported in [14] on the same test set (*ρ* = 0.73), despite that we provide it with a larger training set. This likely reflects that they re-trained the full UniRep model on the stability data, rather than only training the simple downstream model (in their case a single dense layer, in our case gradient boosting). Alternatively, it is possible (though it would be surprising) that the additional training data hurts performance on the prediction task, or that the gradient boosting model was less able to capture the small changes in stability than the dense layer. Future work should evaluate these possibilities, and whether similar gains may be seen by updating the full ESM to solve a particular task.

## 4 Conclusion

Prediction of the effect of mutations on protein stability as well as predicting protein stability itself is an important challenge for molecular and computational biologists. We show that multiple approaches to data-driven modeling of protein stability can be successful, though all depend on the benefits of large, community-accessible datasets. We evaluate these models in a prediction task where they have to extrapolate to a new class of previously-unseen protein, as well as in a task where they have to predict the effects on stability of small changes to a known protein.

We evaluate two kinds of models: relatively small convolutional neural networks trained directly on stability data, and large general-purpose embedding models trained on protein sequences and then interpreted through simple gradient boosting models fit to stability data. Both approaches can solve both classes of problems to a similar degree, with the ESM matching or exceeding the performance of all other individual models. We do not test a third approach, which involves retraining one of the large embedding models to tune it to the task at hand, as it requires substantially greater computational resources. Blending or averaging together multiple models provides a less computationally intensive way of achieving slightly better results. Which approach is best likely reflects the needs and resources of the practitioner.

**Table 1.**
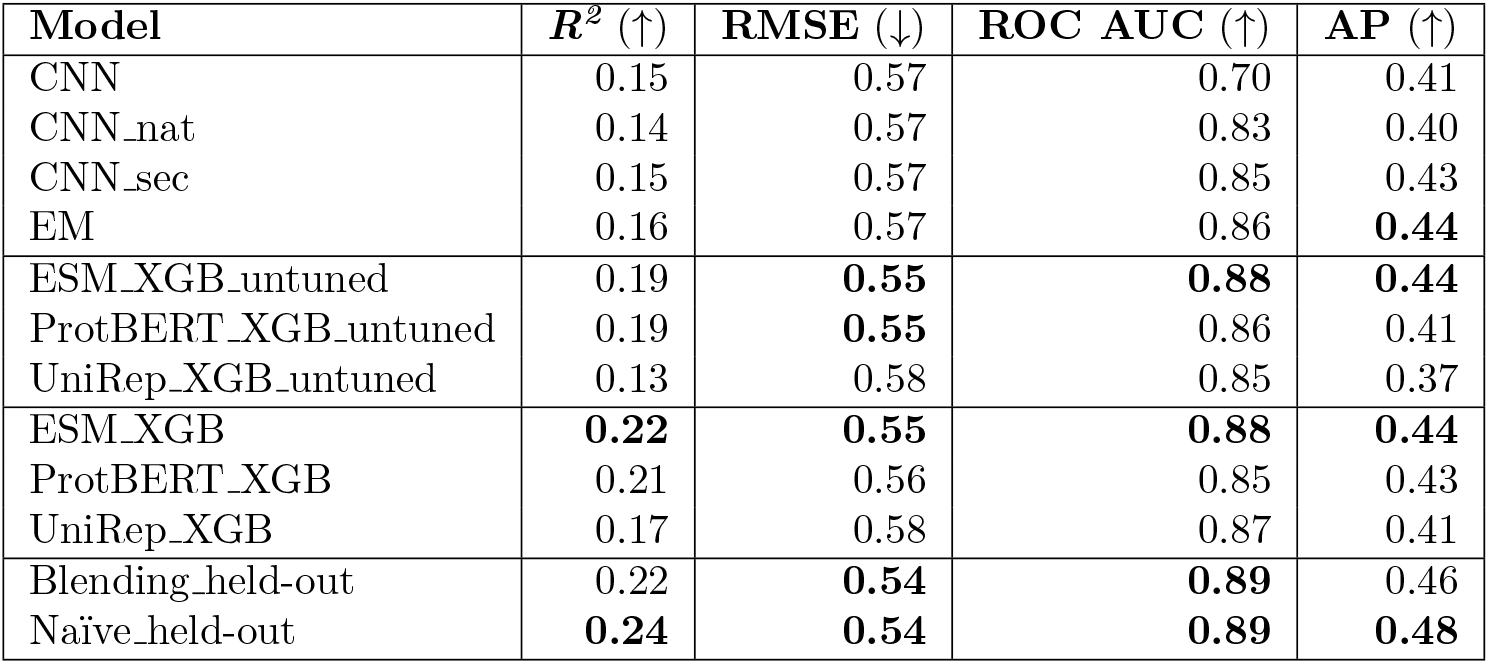
Held-Out Topology Task: Mean Model Performance

**Table 2.** Single-Site Mutation Task: Mean Model Performance

| Model | $R^2$ (↑) | RMSE (↓) | Spearman $\rho$ (↑) |
| --- | --- | --- | --- |
| EM | **0.50** | **0.53** | 0.72 |
| ESM_XGB | **0.50** | **0.53** | **0.73** |
| ProtBERT_XGB | 0.46 | 0.56 | 0.72 |
| UniRep_XGB | 0.41 | 0.58 | 0.67 |
| Blending_ssm | **0.51** | **0.53** | **0.74** |
| Naïve_ssm | **0.51** | **0.53** | **0.74** |

## Acknowledgements

We are thankful for the support and computing resources from the Texas Advanced Computing Center (TACC).

